# Distinct patterns of bioscience doctoral publication disparities by gender and race/ethnicity

**DOI:** 10.1101/2024.10.19.619200

**Authors:** Katie Leap, Gregory S. Payne, Janet S. Sinsheimer, Diana E. Azurdia

**Author notes:** Corresponding author (DEA).

## Abstract

The ability to address the lack of diversity in the Science, Technology, Engineering and Math workforce depends on inclusive and equitable training of doctoral students to succeed in the profession. An important metric used to assess equitable training in bioscience doctoral programs is the number of publications that result from a student’s research. The purpose of this study was to investigate whether there are demographic differences in publication rates among all students in a cohort of bioscience Ph.D. programs at the University of California Los Angeles who graduated between 2011 and 2019. Using institutional data and publication database queries, we determined the number of doctoral publications parsed by authorship position, and the timing of the first publication for each student. The resulting dataset was then analyzed for the relationships between publication categories and student gender, race/ethnicity, and citizenship status. We find that female students published significantly fewer total and co-author papers compared to male students, but had the same number of first-author publications. In contrast, students from underrepresented racial/ethnic groups had fewer first-author papers compared to students from well-represented groups, but similar numbers of total and co-author publications. Publication of the first doctoral paper occurred later for female versus male, and underrepresented versus well-represented students. These results provide evidence for distinct patterns of doctoral publication disparities by gender and race/ethnicity, offering insights into a key metric of bioscience student success and informing potential strategies to achieve equitable outcomes in bioscience doctoral education.

## Introduction

Scientific career success is defined by a variety of metrics, including productivity, the type of independent researcher position obtained, academic impact, length of career, and the ability to attract collaboration [1]. These markers of success develop over the course of years, so they can be less accessible in early career stages. At the doctoral level, productivity, as gauged by publication count and authorship position, is used as an impactful metric during the Ph.D. and for future success. However, a number of studies have uncovered disparities in publication rates linked to gender and racial-ethnic differences at the doctoral level and onwards, with some variation by field [2–15]. As such, ensuring equal access to academic research careers demands more equitable early career publication rates.

In response to these studies and a commitment to addressing hidden academic disparities, Graduate Programs in Bioscience (GPB) at the University of California Los Angeles (UCLA) initiated an examination of publication rate outcomes of graduating students. GPB is a consortium of seven Ph.D. programs with over 500 students in the David Geffen School of Medicine and the College Division of Life Sciences. One mission of GPB is to provide professional development for students and faculty across the consortium, which complements the in-depth research training provided by member programs. As part of this mission, in 2015 GPB began launching various initiatives focused on recruiting and retaining individuals from underrepresented groups (URG) in the biosciences. These initiatives included efforts to enhance culturally inclusive, evidence-based mentorship practices for both faculty and trainees, along with programming aimed at fostering trainee research independence, self-efficacy, leadership skills, and career development. To help guide these efforts, we sought to establish a dataset on publications and time to degree across the GPB programs.

This cohort study investigates potential gender, racial/ethnic, and citizenship differences in doctoral publication numbers and timing of first publication among all GPB graduates from 2011 to 2019. The time period encompasses students who graduated prior to and during the initial implementation phase of GPB initiatives, ending in 2019 to avoid confounding effects of the SARS CoV-2 pandemic. We utilized institutional data and publication database queries with the aim of avoiding potential survey pitfalls such as biased response rates and self-reporting. Here we report doctoral publication number and timing disparities by gender and race/ethnicity, with unexpectedly distinct patterns that differ by authorship position between the two demographic categories.

## Methods

### Study Population

This study was authorized by the UCLA Institutional Review Board, protocol #20-002132. All students who graduated from GPB member programs with a PhD between the years 2011 and 2019 were included in the study. The seven GPB member Ph.D. programs are: Bioinformatics; Human Genetics; Molecular Biology; Molecular, Cellular, & Integrative Physiology; Molecular & Medical Pharmacology; Neuroscience; and Physics & Biology in Medicine. The data span a change in the umbrella program organization in which admissions into four previous programs (Biological Chemistry; Molecular Cell and Developmental Biology; Microbiology, Immunology, and Molecular Genetics; and Cell and Molecular Pathology) were subsumed by the Molecular Biology PhD program. Admissions into all these programs have been grouped under Molecular Biology.

The UCLA Division of Graduate Education (DGE) provided Ph.D. program application data that covered the period from 2002 to 2016 and included demographic and academic information. Gender and ethnicity were self-reported. Because students in the dataset applied to Ph.D. programs during a time when the UCLA application only asked for applicant sex, not yet accommodating non-binary gender affiliations, only self-reported binary designations are available. For this study, we use the term “gender” when referring to these self-reported binary designations for male and female. Based on 2022 application data, in which 2% of applicants identified as non-binary, some students are likely misclassified. For ethnicity, Black, Latine, Native American, Alaskan Native, and Pacific Islander students are categorized as Underrepresented Groups (URG) while White and Asian students are categorized as Overrepresented Groups (ORG). Students who declined to state their ethnicity or who stated their ethnicity as “other” were omitted from the ethnicity aspects of this study. Admission data included citizenship information that was used to define students as international or domestic (U.S. citizen or permanent resident). Academic information included Ph.D. program of study, whether the student was concurrently pursuing an M.D., the doctoral committee chair name, co-chair name if applicable, and matriculation and graduation years. Because of our focus on the early career, we excluded students who had previously obtained an M.D., but retained students who were receiving a concurrent M.D. through the Medical Scientist Training Program program.

### Doctoral Publication Counts

Using the list of student names from the applications submitted to UCLA DGE by all students meeting the study criteria described above, we generated queries to identify publications by the students during the time period of interest. If students had changed their names during the course of their graduate education, this was documented in the record of each name registered by the DGE. Papers published by students were identified by querying PubMed using the R package RISmed [16]. Queries were constructed using student names and the names of the chair of their doctoral committees, including co-chairs if available. For Ph.D. students in the programs we studied, the doctoral committee chair is almost always the student’s faculty research advisor and they generally are included as an author in published doctoral work. To determine the timeframe of doctoral studies, we used matriculation and graduation dates. UCLA operates on a quarter system, with most students starting their Ph.D. programs in October, which is the fall quarter. Due to limitations in PubMed indexing, we were only able to access the calendar year of the publications. Thus, we limited our search to start the calendar year after matriculation and end one calendar year past their graduation year. Any publications during the time frame between matriculation, generally in October, and December of the matriculation year were presumed to be from work done prior to entering the program. We chose one-year post-graduation to capture papers that were submitted or close to completion at graduation. Because there is sometimes a discrepancy between the doctoral committee chair (mentor) of record and the faculty members that a student publishes with, the query was constructed to require either the mentor as an author on the paper or an UCLA affiliation. For example, “BRUIN J[Author] AND (MENTOR D[Author] OR UCLA[Affiliation] OR University of California Los Angeles[Affiliation]) AND 2009[EDAT]: 2017[EDAT]”.

The initial queries identified nearly 10,000 papers, but this number was inflated by false positives arising from students with common names and from papers with large numbers of authors which included those who shared names with the student and their mentor or other UCLA faculty members. Although we searched PubMed using last name and first initial, we curated the article list by requiring a full match for first and last name. If the journal only reported author initials, we required a last name and first initial match, as well as a per-author UCLA affiliation. We eliminated papers that only reported author initials and did not include per-author information on affiliations. We also excluded papers where the student author was listed at a position greater than the 50th author, as in the process of manual checking, we found that these were almost always not the student in question and likely not an important contribution even if a true association. We excluded articles that began with “Correction:” or included the words “withdrawn”, “erratum”, or “corrigendum”. Every student with no articles identified by the query was manually checked to verify the lack of publications. We also manually checked every student with more than 20 articles to verify that there were no false attributions.

One issue with identifying publications by name queries is that accurate publication attribution can be more challenging for women and people of color [17–19]. For students who had changed their names during the course of their doctoral program, we conducted queries using all names provided. Names composed of multiple words can be separated by either spaces or hyphens, but the separating character is not always the same between journals. To address possible complications from hyphenated or multiple-word names, we searched all permutations of multiple-word names to ensure no papers were lost in the query. Some students publish with shortened versions of their first name such that their initials match (e.g. Rob instead of Robert), but the requirement for a full name match missed the variation of the name used in the publication. When students had papers identified by initials but fewer papers identified by a full name match, we verified the name the student used on publications. After all filtering and checking, the search yielded 2924 papers.

Paper counts were calculated by summing the number of all papers identified per student. First-author publications were counted if the student was first in the list of authors. We did not check for co-first author designations when the student is listed second or third with a note that they made an equal contribution to the paper as the first author, because sensitivity analysis showed this was not common in our dataset. Co-author publications were defined as the student having a second or greater position in the author list and were calculated by subtracting the number of first-author publications from the total number of publications.

### Modeling

We modeled three different kinds of outcomes by total publications and by authorship position: the count of papers published by each student, whether or not a student published, or the time to publication or graduation without publication. For count data, we considered the counts to follow a negative binomial distribution. We used both a classic fixed-effects negative binomial regression [20] and a mixed-effects negative binomial regression [21]. Because there are differences in publication rates by scientific field of study, we used the doctoral program of the student as a proxy for their scientific field and fit the program as a random intercept in the model. For the binary outcome of whether a student published or not, we fit a logistic regression [20]. Finally, for the time-to-event data, we used the Kaplan-Meier estimator [22] to model the differences in time to publication and considered students censored if they graduated without publishing. We excluded four students who graduated after 8 years as outliers in this analysis. Figures were made using the R packages ggplot2 and survminer [23,24]. For all statistical tests, p-values are reported and results were considered significant if p < 0.05, but results with p < 0.1 are discussed as notable.

## Results

The seven UCLA bioscience Ph.D. programs listed in Table 1 were selected for analysis based on their membership in a centrally-supported umbrella consortium. These programs provide training across a wide spectrum of bioscience fields. To establish baseline data for the assessment of programmatic initiatives introduced by the consortium, publication rates were determined for 614 students in the member programs who obtained their Ph.D. between 2011 and 2019 (matriculated 2002-2016). Of the students included in the analysis, 10% received their Ph.D. as part of the doctoral training phase of the M.D./Ph.D. program. Table 1 presents the population characteristics by gender, race/ethnicity, citizenship, Ph.D. program, and degree track. The median time to degree for the whole population in the dataset was 5.5 years. There were no differences in time to degree by gender or citizenship status. However, a significant difference was observed by race/ethnicity, with URG students taking longer to graduate (Table 2).

**Table 1.**
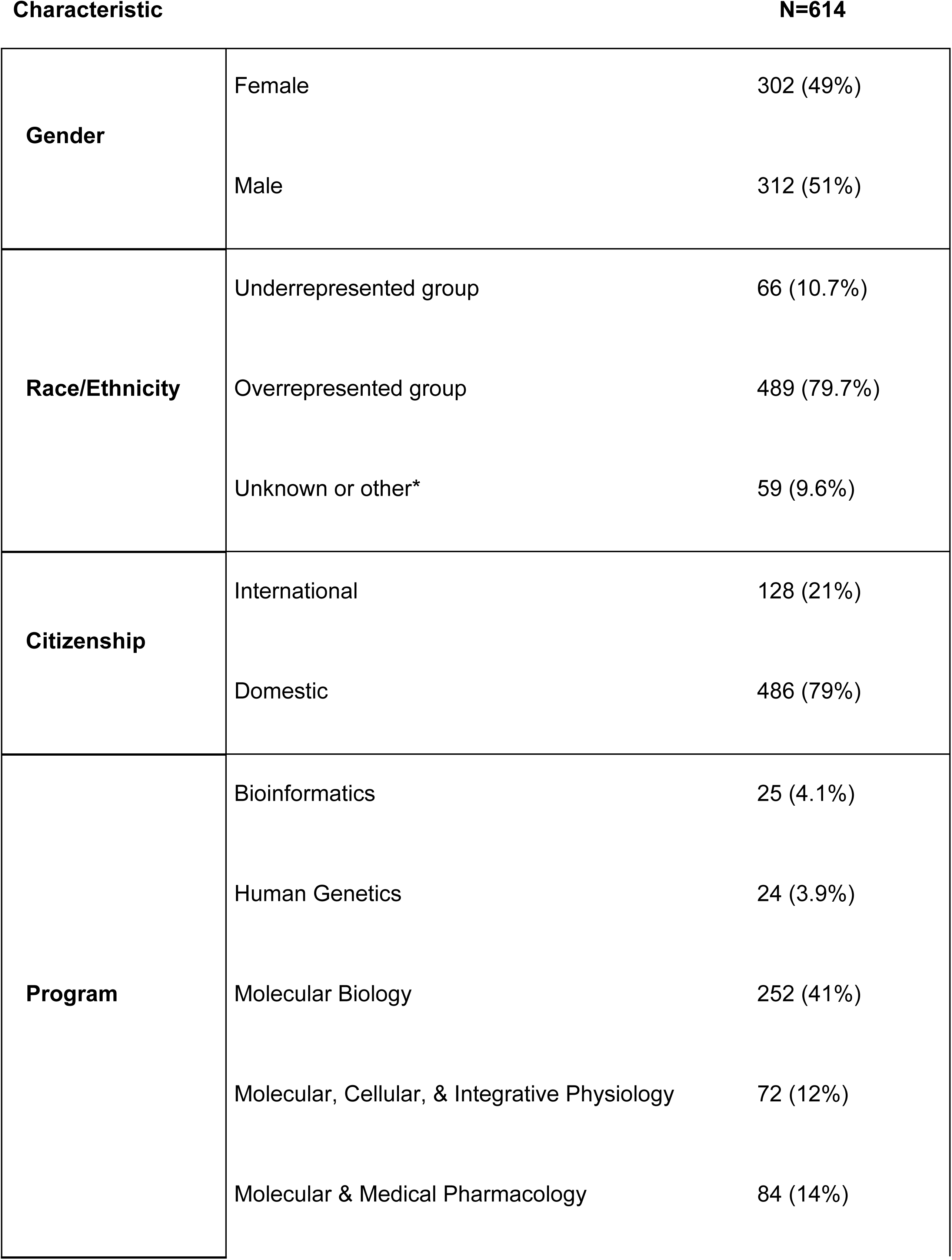
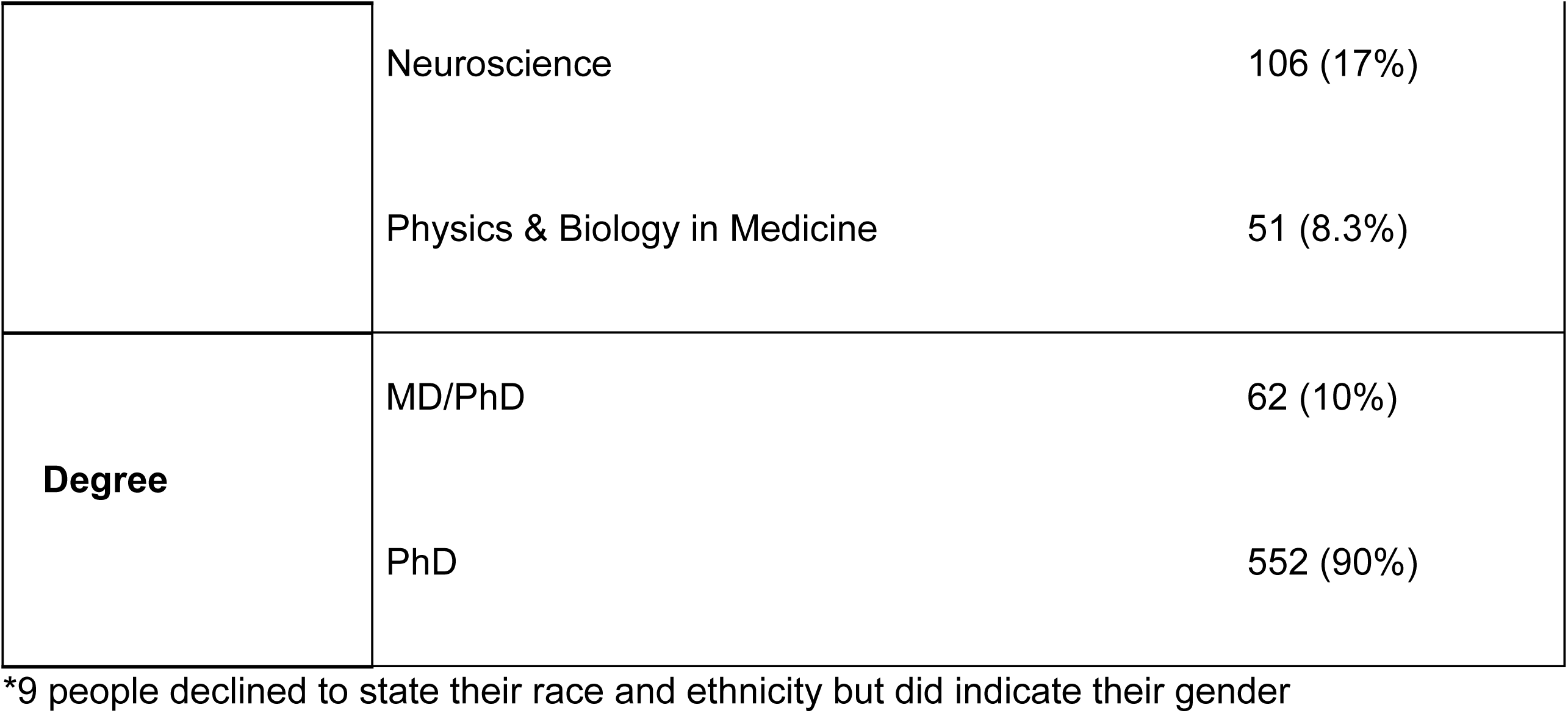
Demographics of study population.

**Table 2.** Disparity in time to degree by race/ethnicity, but not gender or citizenship status.

|  |  | Median (years) | IQR | P-value |
| --- | --- | --- | --- | --- |
| Gender | Female | 5.50 | (4.75, 6.25) | 0.3 |
|  | Male | 5.75 | (4.75, 6.25) |  |
| Race/Ethnicity Group | URG | 5.75 | (5.25, 6.75) | 0.005 |
|  | ORG | 5.50 | (4.75, 6.25) |  |
| Citizenship Status | International | 5.38 | (4.75, 6.25) | 0.7 |
|  | Domestic | 5.75 | (4.75, 6.25) |  |
| Total |  | 5.50 | (4.75, 6.25) |  |

IQR: interquartile range; URG: underrepresented group; ORG: overrepresented group; Domestic: U.S. citizen or permanent resident. P-value determined by the Wilcoxon rank sum test.

To determine the number of doctoral publications for each student we queried the NIH PubMed database for publications between the beginning of the student’s second year in the program and one year after graduation. The query results were then curated to eliminate false attributions, resulting in 2924 papers from 614 students (see Methods). The resulting dataset was analyzed with a focus on comparing the number of publications according to gender, race/ethnicity, and citizenship status. Given the importance of publications as indicators of scholarly productivity in bioscience fields, we first examined whether there were demographic disparities in the percentage of students who failed to publish (Table 3). No significant differences were observed between female and male students (Chi-square test p=0.2), between students in racial/ethnic groups traditionally underrepresented in bioscience (URG: Black, Latine, Native American, Alaskan Native, Native Pacific Islander) and those in groups traditionally overrepresented in bioscience (ORG: White, Asian) (Chi-square test p>0.5), or between international and domestic students (Chi-square test p>0.5).

**Table 3.**
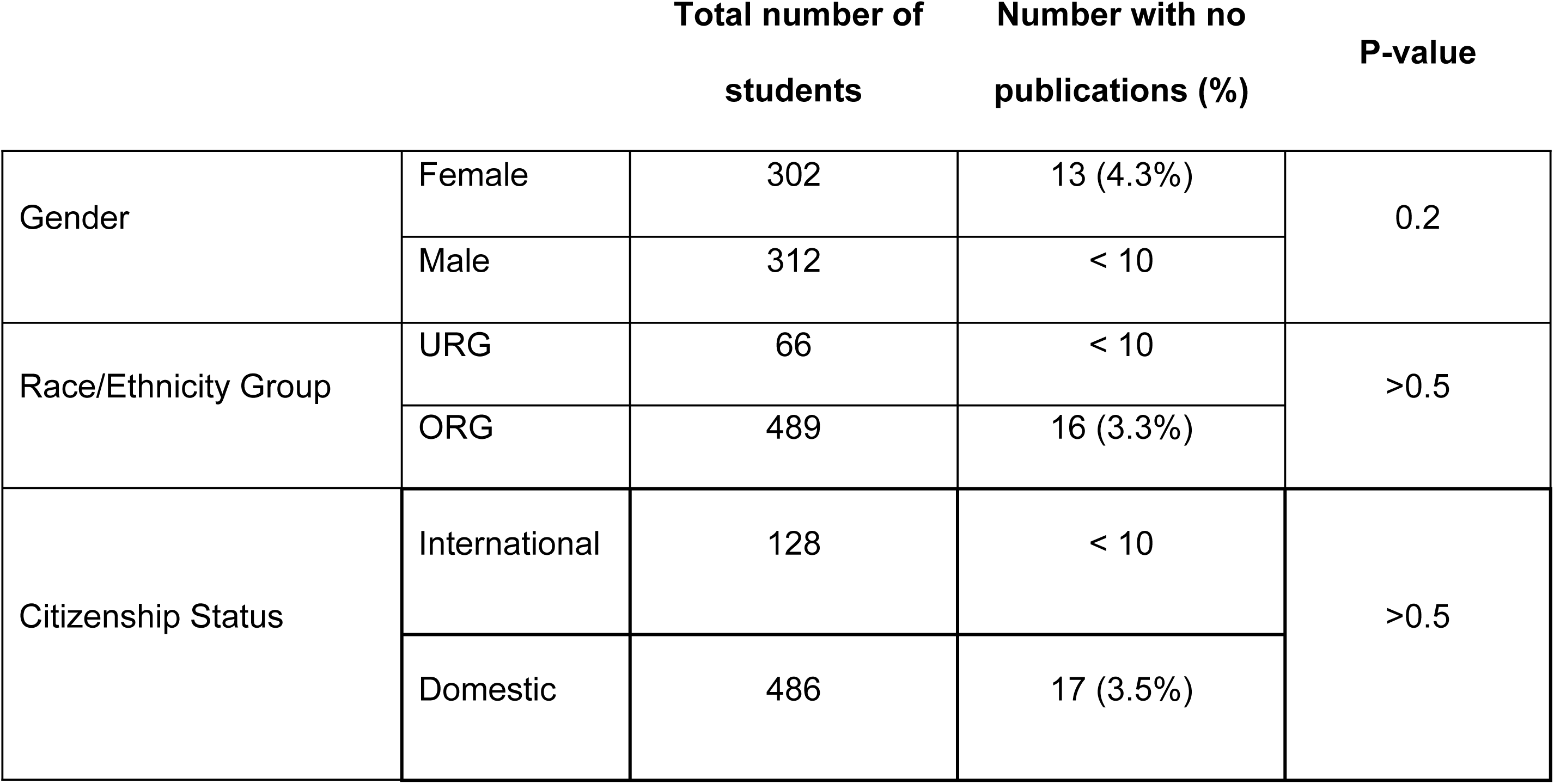
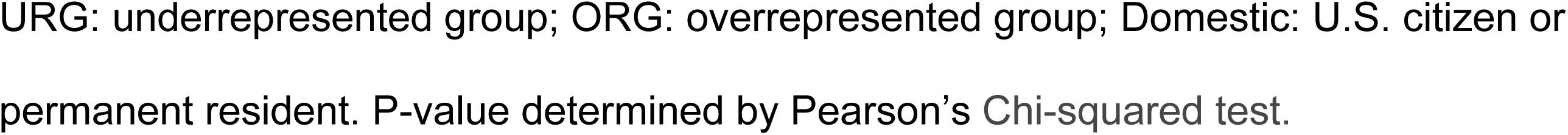
Students with no publications are evenly distributed within gender, race/ethnicity, and citizenship status groups.

|  |  | Total number of students | Number with no publications (%) | P-value |
| --- | --- | --- | --- | --- |
| Gender | Female | 302 | 13 (4.3%) | 0.2 |
|  | Male | 312 | < 10 |  |
| Race/Ethnicity Group | URG | 66 | < 10 | >0.5 |
|  | ORG | 489 | 16 (3.3%) |  |
| Citizenship Status | International | 128 | < 10 | >0.5 |
|  | Domestic | 486 | 17 (3.5%) |  |

URG: underrepresented group; ORG: overrepresented group; Domestic: U.S. citizen or permanent resident. P-value determined by Pearson's Chi-squared test.

We applied the non-parametric unadjusted Wilcoxon Rank Sum Test to compare total publications by student gender or race/ethnicity. Consistent with other reports, female students published significantly less than male students (p<0.001, Table 4A Total paper count, Fig. 1A, Fig. S1A) [9–14]. To probe this disparity in more detail, we compared publication numbers based on the student’s position in the order of authors in each publication. Traditionally in the fields represented in this study, the first author position denotes a more significant contribution than other positions, except for the last position, which is often reserved for the principal investigator who directed the project [25]. Based on this convention, we determined the number of first-author papers for each student and the number of papers where the student was not first-author (co-author papers). Strikingly, there was not a significant difference in the number of first-author publications by female and male students (p=0.13, Table 4A, Fig. 1A, Fig. S1A), but there were significantly fewer co-author publications by female students (p<0.001; Table 4A, Fig. 1A, Fig. S1A). These results suggest that a gender-based difference in co-authorships is a major contributor to the disparity in total publications between female and male students. In contrast to the gender-based results, no significant differences were observed between ORG and URG in total, first-author, or co-author publications (Table 4B, Fig. 1B, Fig. S1B). By citizenship status, there were no significant differences in total and co-author publications, but a reduced number of first-author papers by international students was noted (p=0.063, Table S1).

**Figure 1.**
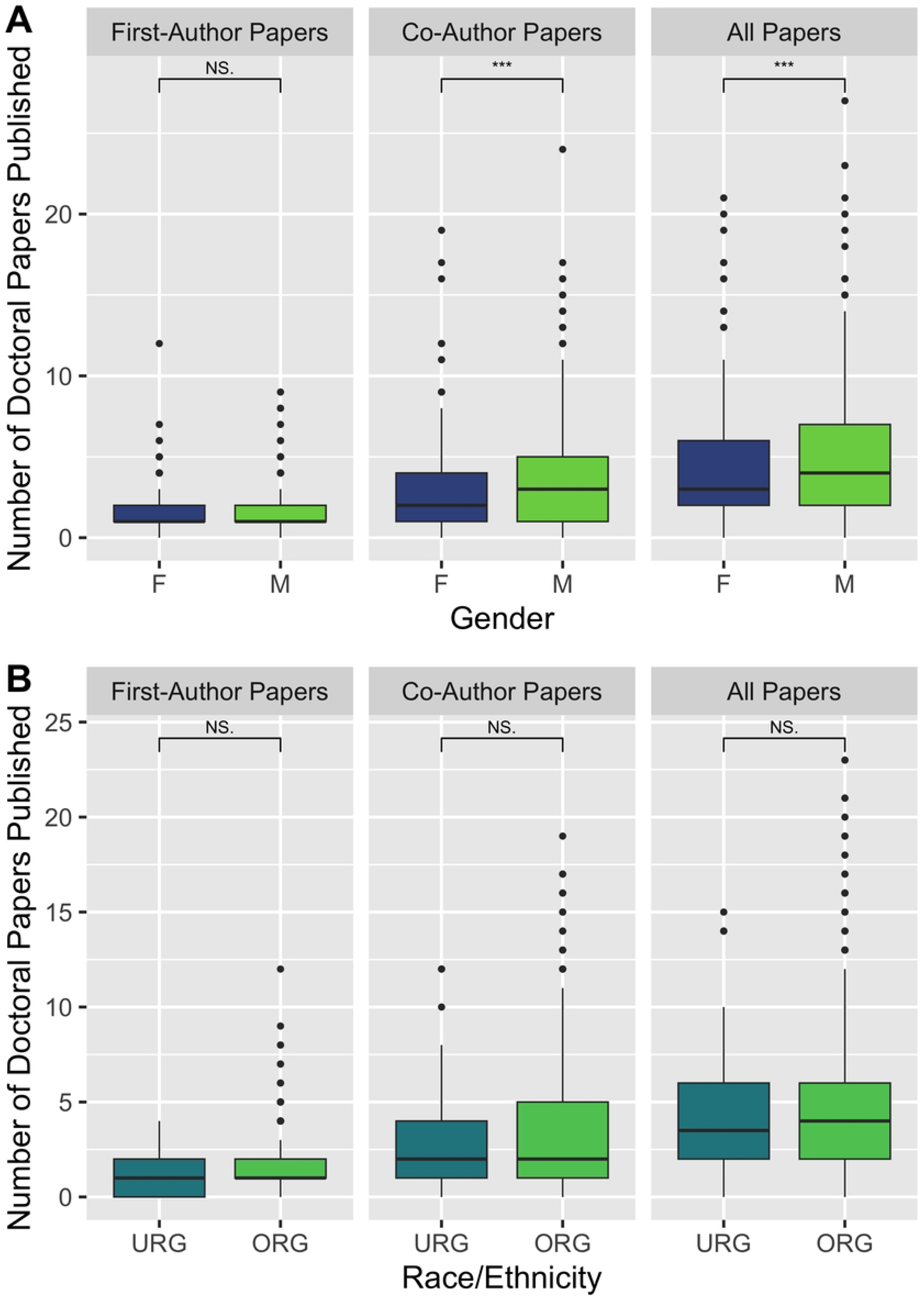
Disparities in unadjusted publications by gender but not race/ethnicity. Distribution of the unadjusted number of doctoral publications by: (A) gender and author position (F: female; M: male); and (B) race/ethnicity and author position (ORG: overrepresented group; URG: underrepresented group). The value for ‘all papers’ is calculated as the sum of the values for first-author papers and co-author papers. The heavy line indicates the median and the boxes indicate the interquartile range. Wilcoxon rank sum test is used to calculate significance. NS: Not significant, P ≥ 0.05; ***: P < 0.001

**Table 4.**
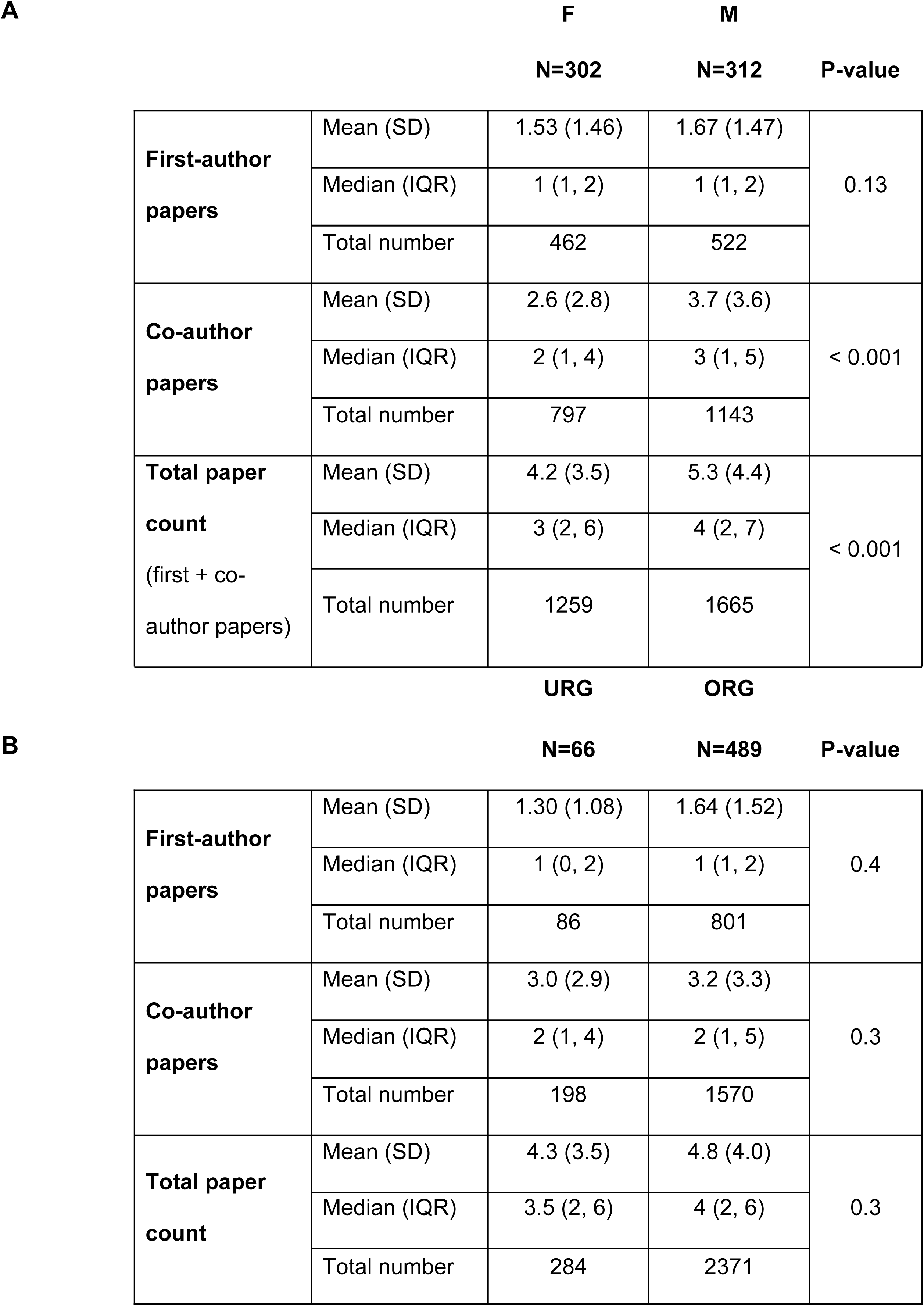
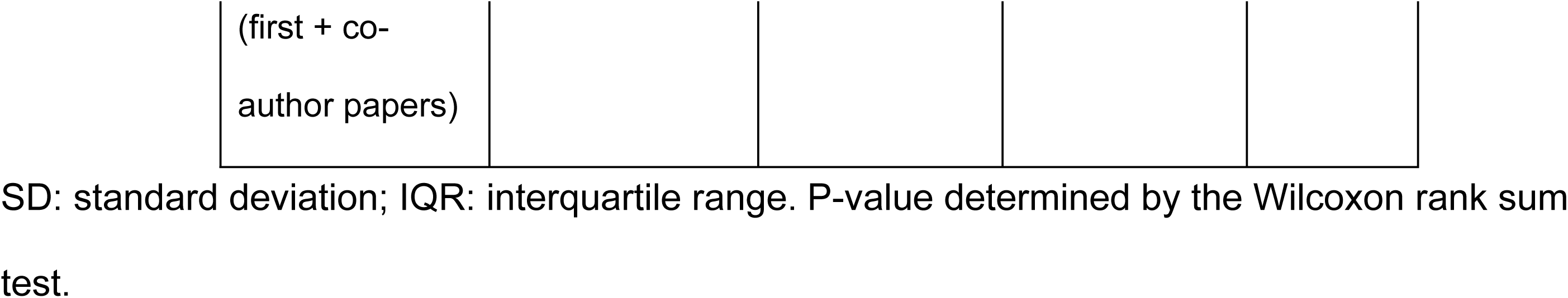
Disparity in unadjusted co-author and total paper counts by gender, but not race/ethnicity.

| **A** |  |  | **F** | **M** | **P-value** |
| --- | --- | --- | --- | --- | --- |
|  |  |  | **N=302** | **N=312** |  |
| **First-author papers** | Mean (SD) |  | 1.53 (1.46) | 1.67 (1.47) | 0.13 |
|  | Median (IQR) |  | 1 (1, 2) | 1 (1, 2) |  |
|  | Total number |  | 462 | 522 |  |
| **Co-author papers** | Mean (SD) |  | 2.6 (2.8) | 3.7 (3.6) | < 0.001 |
|  | Median (IQR) |  | 2 (1, 4) | 3 (1, 5) |  |
|  | Total number |  | 797 | 1143 |  |
| **Total paper count**<br>(first + co-author papers) | Mean (SD) |  | 4.2 (3.5) | 5.3 (4.4) | < 0.001 |
|  | Median (IQR) |  | 3 (2, 6) | 4 (2, 7) |  |
|  | Total number |  | 1259 | 1665 |  |
|  |  |  | **URG** | **ORG** |  |
| **B** |  |  | **N=66** | **N=489** | **P-value** |
| **First-author papers** | Mean (SD) |  | 1.30 (1.08) | 1.64 (1.52) | 0.4 |
|  | Median (IQR) |  | 1 (0, 2) | 1 (1, 2) |  |
|  | Total number |  | 86 | 801 |  |
| **Co-author papers** | Mean (SD) |  | 3.0 (2.9) | 3.2 (3.3) | 0.3 |
|  | Median (IQR) |  | 2 (1, 4) | 2 (1, 5) |  |
|  | Total number |  | 198 | 1570 |  |
| **Total paper count** | Mean (SD) |  | 4.3 (3.5) | 4.8 (4.0) | 0.3 |
|  | Median (IQR) |  | 3.5 (2, 6) | 4 (2, 6) |  |
|  | Total number |  | 284 | 2371 |  |

|  |
| --- |
| (first + co-author papers) |

SD: standard deviation; IQR: interquartile range. P-value determined by the Wilcoxon rank sum test.

To account for the fact that paper counts cannot be below zero or include fractional values, the data were fit to a negative binomial regression model to predict the number of publications by gender, URG, and international status. Furthermore, to factor in possible differences in publication rates between fields of study, we used mixed effects negative binomial regression models that allow for a random effect of Ph.D. program, while fitting gender, URG, and international status as fixed effects. We fit three models predicting numbers of first-author, co-author, and total publications (Table 5).

**Table 5.** Negative binomial mixed-effects regression: gender- and ethnicity/race-based disparities in number of publications.

| Characteristic | First Author Papers |  |  | Co-Author Papers |  |  | Total Papers |  |  |
| --- | --- | --- | --- | --- | --- | --- | --- | --- | --- |
|  | IRR | 95% CI | P | IRR | 95% CI | P | IRR | 95% CI | P |
| Female | 0.96 | 0.83, 1.11 | 0.6 | 0.75 | 0.64, 0.88 | <0.001 | 0.82 | 0.72, 0.93 | 0.002 |
| URG | 0.78 | 0.61, 1.0 | 0.046 | 1.01 | 0.79, 1.30 | >0.9 | 0.93 | 0.76, 1.13 | 0.4 |
| International | 0.91 | 0.75, 1.10 | 0.3 | 1.09 | 0.90, 1.33 | 0.4 | 1.02 | 0.87, 1.2 | 0.8 |

Coefficients from negative binomial mixed effects regression, incorporating fixed effects of gender (male as reference), race/ethnicity (ORG: overrepresented group as reference), and international status (domestic as reference) and a random effect of doctoral program. URG: underrepresented group. IRR = Incidence Rate Ratio, CI = Confidence Interval, P = p-value

The results of this approach also provide evidence for gender-based disparities in total and co-author publications: the total publication incidence rate ratio (IRR) for female versus male students was 0.82 (p=0.002), co-author IRR was 0.75 (p<0.001), first author IRR was 0.96 (p=0.6). That is, as an example, the data predict that for every ten papers published by a male student, a female student publishes only 8 papers. In contrast, we observed that URG status was significantly associated with a negative effect on first-author publications (IRR 0.78, p=0.046), but not on co-author (IRR 1.01, p>0.9) or total (IRR 0.93, p=0.4) publications. These findings reveal a race/ethnicity-based disparity in first-author publications. There were no observed differences between international and domestic students for any of the publication categories. Similar results for each demographic category were obtained using fixed effects negative binomial regression models without accounting for possible publication differences by field (Table S2).

The negative binomial regression analyses uncovered distinct patterns of publication number disparities between gender and race/ethnicity categories. In light of these findings, we explored if there were differences in whether students publish at all by logistic regression, accounting for the effects of gender, race/ethnicity, and citizenship status on whether a student had no publications in any of the authorship categories (Table 6). Although the patterns of female/male- or URG/ORG-based differences across authorship categories by logistic regression parallel those in the negative binomial analyses, only the gender disparity in co-author papers was significant, with female students having virtually half the odds of publishing any co-author paper compared to male students. Based on this analysis, URG students had notably lower odds of any first-author publications compared to ORG students. International students were no more or less likely to publish overall compared to domestic students but had decreased odds of any first-author papers (OR = 0.59; p = 0.026) yet increased odds of any co-author papers (OR = 1.83; p = 0.06).

**Table 6.** Logistic regression: disparities in odds of publishing by gender and international status.

|  | First Author Papers |  |  | Co-Author Papers |  |  | Total Papers |  |  |
| --- | --- | --- | --- | --- | --- | --- | --- | --- | --- |
| Characteristic | OR | 95% CI | P | OR | 95% CI | P | OR | 95% CI | P |
| Female | 0.91 | 0.60, 1.37 | 0.6 | 0.47 | 0.29, 0.77 | 0.003 | 0.53 | 0.19, 1.36 | 0.2 |
| URG | 0.59 | 0.33, 1.07 | 0.074 | 1.11 | 0.56, 2.44 | 0.8 | 0.80 | 0.25, 3.54 | 0.7 |
| International | 0.59 | 0.37, 0.95 | 0.026 | 1.83 | 0.99, 3.62 | 0.066 | 1.64 | 0.52, 7.24 | 0.4 |

OR = Odds Ratio, CI = Confidence Interval, P = p-value; Male as reference for gender, ORG as reference for race/ethnicity, domestic as reference for international.

The timing of a student’s initial doctoral publication in any category of authorship has the potential to impact both the total number of graduate publications and other factors that can contribute to student success during their graduate studies (and onward in their career). For example, publications are commonly used as metrics for graduate fellowships, appointments to federally-funded predoctoral training programs, funding for travel and registration to attend conferences, and other types of awards and distinctions. Accordingly, we used the Kaplan-Meier method to compare the timing of the initial graduate publication by female versus male students [26]. As presented in Fig. 2A, the probability of not having published a paper declined for female students more slowly than male students (p=0.05), with a gap becoming evident by the end of year 2 and dissipating by the end of year 6. Thus, there was a notable delay in the time to first graduate school publication for female students compared to male students. When the timing of initial publication was analyzed by authorship position, the data revealed that students generally published their initial first-author paper later than a co-author paper (Figures 2B and 2C). Notably, the timing of the initial first-author paper was not significantly different between female and male students (p=0.39; Figure 2B), whereas a significant delay was evident in the timing of the initial co-author publication by female compared to male students (p=0.0047; Figure 2C). These findings provide evidence of a gender-based disparity in the timing of a student’s initial publication in graduate school, which can be largely attributed to a difference in the timing of the initial co-author publication. A significant delay in the timing of initial publication by URG compared to ORG was also apparent by Kaplan-Meier analysis (Figure 3A). When analyzed by authorship position, differences in the timing of initial publication were not significant, likely due to the small number of URG students (Figure 3B-C).

**Figure 2.**
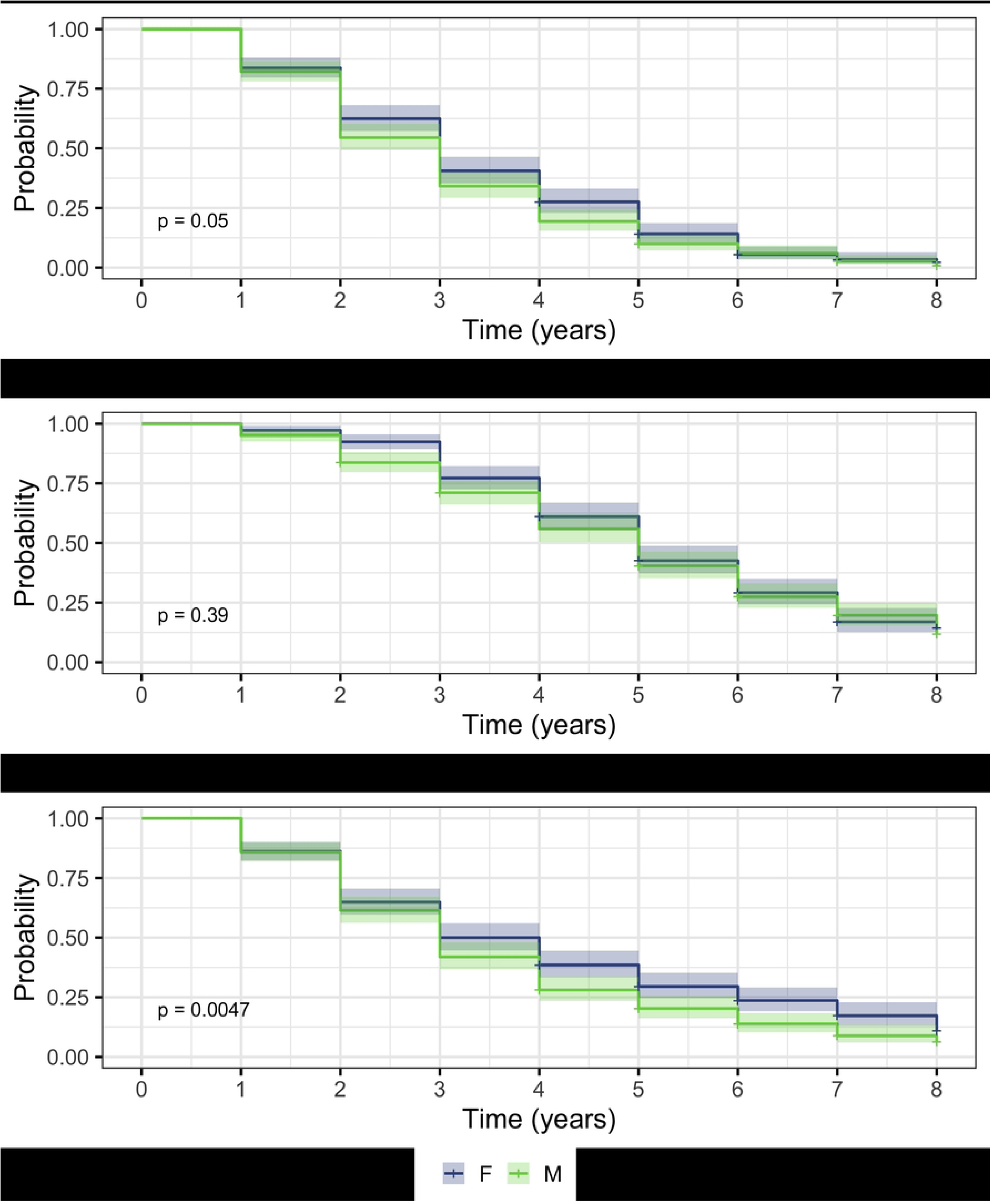
Disparity in time to first graduate co-author publication by gender. Time to first publication by author position represented by Kaplan-Meier curves. The probability that a student remains in the program without a publication is presented as a function of time spent in the program, with the shaded area representing the 95% confidence interval. Students who graduated with no publications of a given authorship type are considered to be censored in the dataset, which is represented by a + symbol. Four students who graduated after 8 years were excluded as outliers. (A) Probability that a student has not published as a function of years spent in their graduate program. (B) Probability that a student has not published a first-author paper as a function of years spent in their graduate program. (C) Probability that a student has not published a co-authored paper as a function of years spent in their graduate program.

**Figure 3.**
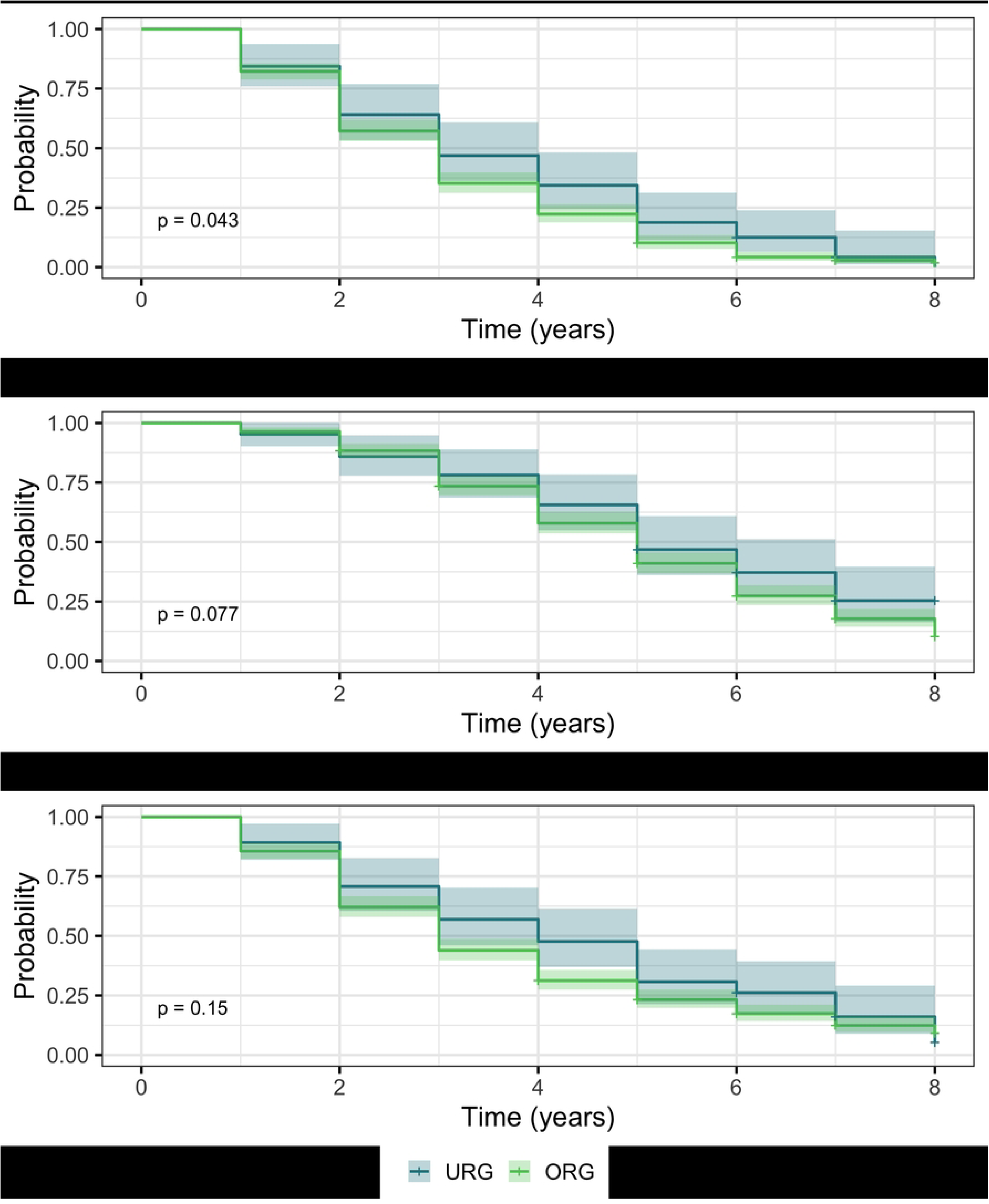
Disparity in time to first graduate publication by race/ethnicity. Time to first publication by author position represented by Kaplan-Meier curves. The probability that a student remains in the program without a publication is presented as a function of the time spent in the program, with the shaded area representing the 95% confidence interval. Students who graduated with no publications of a given authorship type are considered to be censored in the dataset, which is represented by a + symbol. Four students who graduated after 8 years were excluded as outliers. (A) Probability that a student has not published as a function of years spent in their graduate program. (B) Probability that a student has not published a first-author paper as a function of years spent in their graduate program. (C) Probability that a student has not published a co-authored paper as a function of years spent in their graduate program.

## Discussion

Publications serve as important indicators of research potential during Science, Technology, Engineering, and Math (STEM) education and are broadly used as measures of success in STEM careers. Consequently, inequities that arise at early stages of training can have long-range impacts. In this study, we assembled a dataset of doctoral publications by students in a consortium of bioscience Ph.D. programs and analyzed the dataset for publication number and timing disparities across different demographic categories. The results provide evidence for distinct patterns of disparities by gender and race/ethnicity, distinguished by authorship position. These findings offer insights that can inform strategies to reduce inequities that arise during doctoral research training and extend into later stages of academic careers [27,28].

Our analyses by gender extend existing documentation that women publish less than men in STEM fields [3–7,15], and demonstrate that this difference is already apparent at the Ph.D. level in a cohort of bioscience students, consistent with other reports that publication disparities arise at this early career stage [9–14]. Notably, our findings also highlight underappreciated aspects of publication gender bias. By including authorship position in our studies, the data reveal that the diminished number of total publications by female students is driven by fewer co-authored publications. Furthermore, we find that female students publish their initial doctoral paper later than male students, and the initial paper for both genders is more likely to be a co-authorship. Together, these results suggest that co-authorship may be a key early determinant of gender-based disparities in STEM careers.

A slower time for women to publish their initial doctoral paper, regardless of authorship position, may have potentially significant negative implications for their graduate education. The absence of a doctoral publication can impact not only tangible outcomes like fellowships and prizes but also less tangible aspects of professional development such as self-efficacy. Self-efficacy, an individual’s belief in their capacity to perform specific tasks [29], is associated with higher research productivity, in addition to increased measures of personal satisfaction and resilience [30–32]. As such, self-efficacy is an important psychosocial component of doctoral student professional development. Considering that publications are a significant metric of success, the initial doctoral paper serves as a strong indicator of performance accomplishments, and a key source of self-efficacy [33]. By this reasoning, the gender disparity in the time to initial publication disadvantages female students by delaying a substantial boost in self-efficacy and associated benefits.

In the bioscience fields represented in this study, co-authorship (i.e. author position between first and last authors) can be interpreted as an indicator of collaboration in a project, based on the academic convention of assigning first and last authorship to the project leaders. From the perspective of collaboration, a number of factors may contribute to co-authorship gender disparity. For example, there may be a bias toward women in conferring authorship for collaborative contributions, as reported by Ross et al. [34]. Other possibilities include ineffective mentorship and/or fewer or delayed opportunities for women to participate in collaborations [35].

An intervention to address gender-based disparities in co-authored publications and time to initial paper would be to increase awareness among mentors and mentees. Such conversations could be introduced into mentorship training and development activities for faculty and trainees, as well as in other forums aimed at promoting trainee success. Raising awareness could be coupled with concerted encouragement of mentors to create early opportunities for collaboration, increasing the chances for co-authorship and thereby shortening the time to initial publication. Collaboration could also present students with experiences that are known to be sources of self-efficacy in addition to mastery of skills, for example: the ability to vicariously learn, receive positive feedback that fosters growth, and experience excitement through contributing to a larger project [33,36,37]. Optimally, access to teamwork and collaborations that lead to co-authorship would be an integral part of doctoral training in the biosciences, with attention to delegating opportunities equitably.

Our regression analyses revealed a pattern of disparity by race/ethnicity that is distinct from that of the gender-based disparities. In the case of publication numbers, when compared to ORG students, URG students had fewer first-author papers but similar numbers of co-author and total papers. In the case of time to initial publication, there was a delay for URG students which mirrored that observed between the genders.

Even without a significant difference in total publications, the race/ethnicity disparity in the number of first-author publications has the potential to be especially impactful in academic settings, given that the first-author position normally reflects leadership of a project and often is considered a more important indicator of independent scholarship than co-authorship. A possible example from our data consistent with this view is an effect on time to degree. We observed that the first-author publication disparity was associated with a slower time to doctoral degree for URG students. In contrast, there was no difference in time to degree by gender, where women had the same number of first-author publications as men, although experiencing a disparity in time to initial publication comparable to that of URG students.

It is worth noting that the race/ethnicity results were more nuanced than those from the gender analysis: the race/ethnicity disparity in the number of first-author publications was apparent only in regression models, not in unadjusted analysis, and it was not possible to extend the difference in timing of initial publication to authorship position. We attribute these limitations to effects on the statistical power of our analyses that stem from the small number of underrepresented students in the cohort. This is likely to be less of an issue in follow-up studies because the diversity across the cohort programs has more than doubled in recent years.

We propose the mentoring relationship between student and faculty research adviser as a target to address the race/ethnicity-based publication disparities. Effective mentorship plays a crucial role in the ability of mentees to persist and succeed in research careers, profoundly influencing their research productivity, self-efficacy, and ultimate career fulfillment [38]. However, race/ethnicity is a factor that can impede effective mentorship: racial/ethnic disparities in access to high-quality mentorship have been suggested as barriers to publishing and funding [39,40]. Moreover, issues arising from cultural diversity, such as race and ethnicity, can hinder the continued engagement of URG mentees with their mentors [41]. These findings suggest that efforts to reduce the race/ethnicity-based publication inequities would benefit from an early mentorship intervention that supports cultural awareness and responsiveness. An example would be Culturally Aware Mentorship (CAM), an evidence-based training curriculum designed to facilitate mentors’ understanding of their intrapersonal views on race and ethnicity and how those views may then influence interpersonal interactions, ultimately bringing awareness of how their biases and privileges may play out in their mentoring relationships [42–45]. As part of CAM, mentors could be made aware of publication disparities and encouraged to consider the potential impact of their biases on student project and authorship decisions. Supplemental interventions could include providing professional development venues for trainees to obtain secondary mentorship or access to broader networks of support that aim to address disparities in doctoral training and later career stages.

The patterns of publication disparities defined in our study derive from an analysis of a subset of Bioscience Ph.D. programs in the UCLA School of Medicine and Division of Life Sciences. This subset covers a wide spectrum of biomedical and life science areas, from Medical Informatics to Molecular Biology, suggesting that the results are broadly applicable. However, additional studies are needed to assess the full scope of these types of doctoral publication inequities across STEM programs both within and beyond UCLA.

With the specific demographic disparities uncovered in this baseline study, it will be important to monitor the effects of consortium initiatives aimed at reducing inequity in subsequent student cohorts. Follow-up research on student publications should be coupled with surveys to probe student experiences in domains that include mentorship, program structure, and psychosocial development [46]. Ultimately, the goal is to inform evidence-based interventions that lead to fully inclusive doctoral programs and career outcomes in STEM.

## Acknowledgments

This work was inspired by discussions at Howard Hughes Medical Institute Science Education Department with David Asai and Clif Poodry. We thank Hua Zhou, Esteban Dell’Angelica, and Sylvia Hurtado for insightful conversations and comments on the manuscript and Raquel Aragón and Melissa J. Spencer for supporting this work. We are grateful to the UCLA Division of Graduate Education for providing data. This manuscript is dedicated to the memory of our coauthor Janet Sinsheimer.

## Supporting Information

**S1 Figure: Distribution of paper counts by gender and race/ethnicity**

Distribution of the percentage of students by publication count (A) by student gender (B) by student race/ethnicity. Six students who had more than 20 papers were removed for the purposes of legibility.

**S1 Table: No significant disparity in total and authorship position paper counts by citizenship.**

SD: standard deviation; IQR: interquartile range, Domestic: U.S. citizen or permanent resident. p-value determined by the Wilcoxon rank sum test

**S2 Table: Negative binomial regression fixed effects model is similar to the mixed effects model**

Coefficients from negative binomial regression incorporating the effects of gender (male as reference), race/ethnicity (overrepresented group as reference) and international status (domestic as reference). URG: underrepresented group. IRR = Incidence Rate Ratio, CI = Confidence Interval, P = p-value

## Notes

### Competing Interest Statement

The authors have declared no competing interest.

### Summary of Updates

The revision includes supplemental materials which were not included in the original submission. The author order was also corrected.

