## Supplemental Tables and Figures for "Distinct patterns of bioscience doctoral publication disparities by gender and race/ethnicity"

### Supporting Information

#### **S1 Table: No significant disparity in total and authorship position paper counts by citizenship.**

|  |  | **International** | **Domestic** |  |
| --- | --- | --- | --- | --- |
|  |  | **N=128** | **N=486** | **P value** |
| **First author papers** | Mean (SD) | 1.52 (1.66) | 1.63 (1.41) | 0.063 |
|  | Median (IQR) | 1 (0, 2) | 1 (1, 2) |  |
|  | Total number | 194 | 790 |  |
| **Co-author papers** | Mean (SD) | 3.3 (3.4) | 3.1 (3.3) | 0.5 |
|  | Median (IQR) | 2 (1, 5) | 2 (1, 4) |  |
|  | Total number | 423 | 1517 |  |
| **Total paper count**  (first + coauthor papers) | Mean (SD) | 4.8 (4.3) | 4.7 (3.9) | 0.8 |
|  | Median (IQR) | 3.5 (2, 7) | 4 (2, 6) |  |
|  | Total number | 617 | 2307 |  |

SD: standard deviation; IQR: interquartile range, Domestic: U.S. citizen or permanent resident. p-value determined by the Wilcoxon rank sum test

#### **S2 Table: Negative binomial regression fixed effects model is similar to the mixed effects model**

|  | **First Author Papers** | | | **Co-Author Papers** | | | **Total Papers** | | |
| --- | --- | --- | --- | --- | --- | --- | --- | --- | --- |
| **Characteristic** | **IRR** | **95% CI** | **P** | **IRR** | **95% CI** | **P** | **IRR** | **95% CI** | **P** |
| Female | 0.94 | 0.81, 1.09 | 0.4 | 0.73 | 0.62, 0.86 | <0.001 | 0.80 | 0.70, 0.91 | <0.001 |
| URG | 0.78 | 0.61, 1.0 | 0.055 | 0.97 | 0.76, 1.25 | 0.8 | 0.90 | 0.74, 1.10 | 0.3 |
| International | 0.90 | 0.74, 1.08 | 0.2 | 1.11 | 0.92, 1.36 | 0.3 | 1.03 | 0.88, 1.21 | 0.7 |

Coefficients from negative binomial regression incorporating the effects of gender (male as reference), race/ethnicity (overrepresented group as reference) and international status (domestic as reference). URG: underrepresented group. IRR = Incidence Rate Ratio, CI = Confidence Interval, P = p-value

#### **S1 Figure: Distribution of paper counts by gender and race/ethnicity**

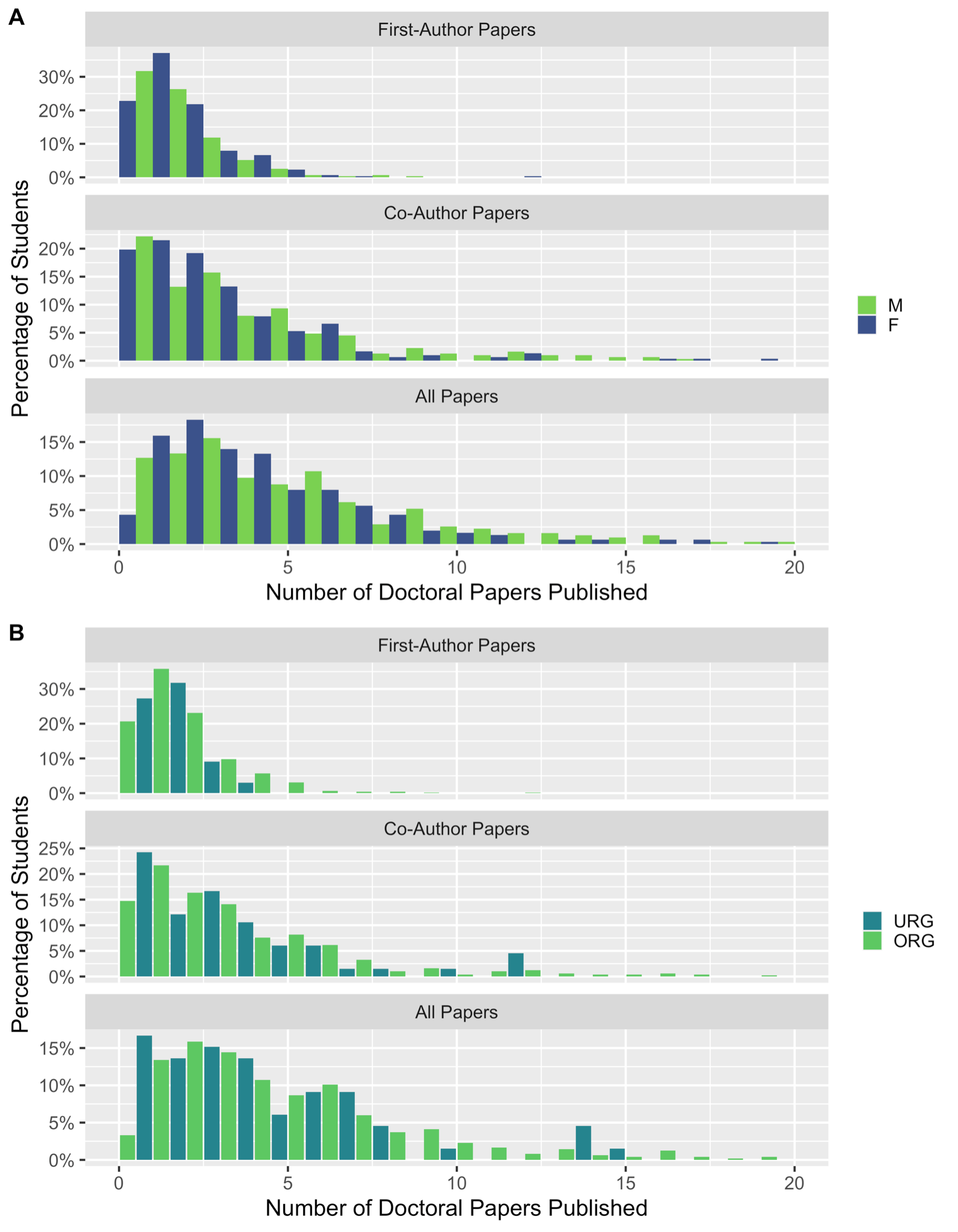

Distribution of the percentage of students by publication count (A) by student gender (B) by student race/ethnicity. Six students who had more than 20 papers were removed for the purposes of legibility.
